# A direct RNA-protein interaction atlas of the SARS-CoV-2 RNA in infected human cells

**DOI:** 10.1101/2020.07.15.204404

**Authors:** Nora Schmidt, Caleb A. Lareau, Hasmik Keshishian, Randy Melanson, Matthias Zimmer, Luisa Kirschner, Jens Ade, Simone Werner, Neva Caliskan, Eric S. Lander, Jörg Vogel, Steven A. Carr, Jochen Bodem, Mathias Munschauer

## Abstract

SARS-CoV-2 infections pose a global threat to human health and an unprecedented research challenge. Among the most urgent tasks is obtaining a detailed understanding of the molecular interactions that facilitate viral replication or contribute to host defense mechanisms in infected cells. While SARS-CoV-2 co-opts cellular factors for viral translation and genome replication, a comprehensive map of the host cell proteome in direct contact with viral RNA has not been elucidated. Here, we use RNA antisense purification and mass spectrometry (RAP-MS) to obtain an unbiased and quantitative picture of the human proteome that directly binds the SARS-CoV-2 RNA in infected human cells. We discover known host factors required for coronavirus replication, regulators of RNA metabolism and host defense pathways, along with dozens of potential drug targets among direct SARS-CoV-2 binders. We further integrate the SARS-CoV-2 RNA interactome with proteome dynamics induced by viral infection, linking interactome proteins to the emerging biology of SARS-CoV-2 infections. Validating RAP-MS, we show that CNBP, a regulator of proinflammatory cytokines, directly engages the SARS-CoV-2 RNA. Supporting the functional relevance of identified interactors, we show that the interferon-induced protein RYDEN suppresses SARS-CoV-2 ribosomal frameshifting and demonstrate that inhibition of SARS-CoV-2-bound proteins is sufficient to manipulate viral replication. The SARS-CoV-2 RNA interactome provides an unprecedented molecular perspective on SARS-CoV-2 infections and enables the systematic dissection of host dependency factors and host defense strategies, a crucial prerequisite for designing novel therapeutic strategies.

## INTRODUCTION

At the end of 2019, the rapid spread of a novel severe acute respiratory syndrome-related coronavirus (SARS-CoV-2) around the globe has led to a worldwide spike in a SARS-like respiratory illness termed Coronavirus Disease 2019 (COVID-19)^1^. Due to the absence of effective antiviral therapy, COVID-19 has taken hundreds of thousands of lives to date and resulted in unprecedented socioeconomic disruptions.

A prerequisite for understanding SARS-CoV-2 infections and enabling novel therapeutic strategies is obtaining a detailed map of the molecular events and perturbations occurring as SARS-CoV-2 infects human host cells. SARS-CoV-2 is an enveloped, positive-sense, single-stranded RNA virus that, upon infection of a host cell, deploys a ‘translation-ready’ RNA molecule, which engages the protein synthesis machinery of the host in order to express a limited number of viral proteins critical for its replication^2^. Thus, similar to other RNA viruses, SARS-CoV-2 is inherently dependent on recruiting host cell factors and machinery, including regulators of RNA stability, localization, and translation, to facilitate virus replication and the production of viral progeny. For the host cell, on the other hand, it is crucial to detect the presence of a viral pathogen and activate appropriate innate immune response pathways^3,4^. To understand this interplay between virus and host, it is essential to characterize with molecular detail which host proteins make direct contact with viral RNA and may function as host dependency factors or antiviral regulators. To date, studies on SARS-CoV-2 infected human cells have primarily focused on characterizing expression or modification changes in the host cell transcriptome^5–7^ or proteome^8–10^. While several studies described protein-protein interactions of recombinantly expressed viral proteins in uninfected cells^10,11^, no study has comprehensively identified direct interactions between viral RNA and the host cell proteome in infected human cells.

To improve our understanding of the host factors that contribute to the regulation of SARS-CoV-2 RNA during infection, we sought to obtain an unbiased picture of the cellular proteins that directly bind to the SARS-CoV-2 RNA in infected human cells. Recent advances in RNA capture and quantitative mass spectrometry approaches^12–14^ have made this endeavor highly tractable. Among the available technologies, we focused on those that use ultraviolet (UV) crosslinking to create covalent bonds between RNA and directly bound proteins, as opposed to chemical crosslinkers that also stabilize indirect interactions^15^. Further, we wanted to ensure that the identified proteins bind directly to the SARS-CoV-2 RNA, excluding approaches that assess differential interactions across all cellular RNAs in response to infection. To satisfy these requirements, we selected RNA antisense purification and quantitative mass spectrometry (RAP-MS)^12,13^, which implements a denaturing purification procedure to capture and identify proteins that crosslink directly to the SARS-CoV-2 RNA.

Using RAP-MS, we globally identify host cell proteins that directly bind to the SARS-CoV-2 RNA in infected human cells and integrate these binding events with proteome abundance changes induced by viral infection. Our work highlights the molecular interactions that underlie both virus replication and host defense mechanisms and further establishes that therapeutic inhibition of direct RNA binders can modulate SARS-CoV-2 replication.

## RESULTS

### Capturing the SARS-CoV-2 RNA and its protein binding partners in human cells

To purify the SARS-CoV-2 RNA and the complement of directly crosslinked cellular proteins from virally infected human cells, we designed a pool of biotinylated DNA oligonucleotides antisense to the positive-sense SARS-CoV-2 RNA, such that one probe binding site occurs roughly every 400 bases in the ~30 kb SARS-CoV-2 genome. As a cellular system, we selected the human HuH-7 cell line. In addition to being permissive to both SARS-CoV-1 and SARS-CoV-2 replication^16,17^, the cellular proteins bound to all polyadenylated RNAs (poly(A)-RNA) have been identified in these cells^18^. Further, a previous study employed chemical crosslinking and RNA antisense purification in a HuH-7-derived cell line to identify proteins that directly or indirectly associate with the genomic RNAs of Dengue and Zika viruses^19^, two unrelated positive-sense RNA viruses.

To test if our pool of antisense capture probes was suitable for the purification of the SARS-CoV-2 RNA from infected HuH-7 cells, we performed RAP-MS 24 hours after infection. Expanding upon the established RAP-MS procedure^13^, we implemented a covalent protein capture step following the release of SARS-CoV-2-bound proteins, which allowed us to identify the RNA sequences crosslinked to purified proteins (Figure 1a, see Methods). While this approach yielded near-complete sequence coverage of the SARS-CoV-2 genome in RNA antisense purifications, most of the signal was observed near probe binding sites (Supplementary Figure 1a). Importantly, sequencing reads originating from SARS-CoV-2 RNA made up 93% and 92% of all mapped reads in two replicate experiments (Supplementary Figure 1b).

**Figure 1.**
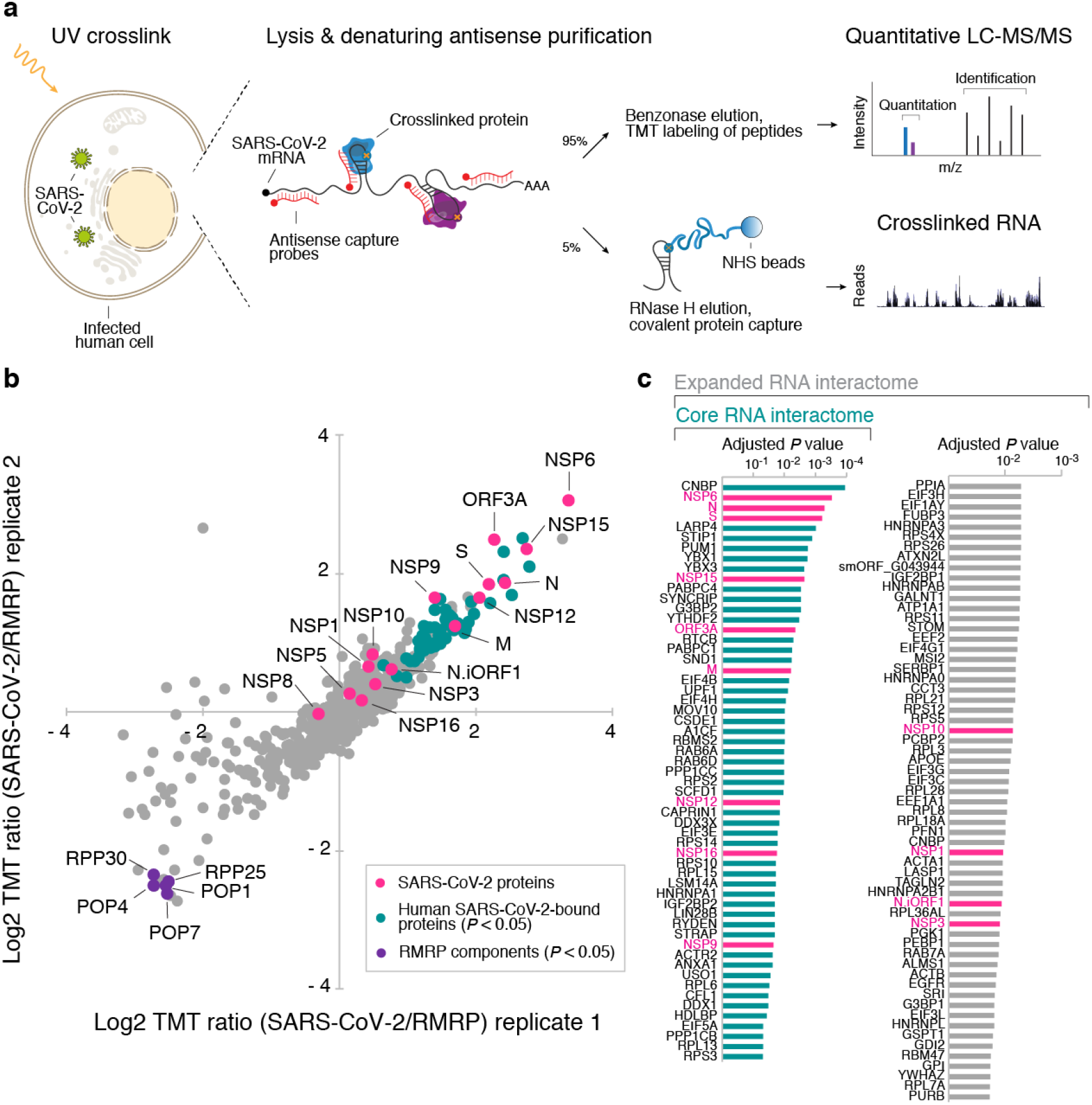
RAP-MS captures the direct RNA-protein interactome of the SARS-CoV-2 RNA in infected human cells. **a**, Outline of the RAP-MS method to identify proteins bound to SARS-CoV-2 RNA and their crosslinked RNA sequences. **b**, Quantification of SARS-CoV-2 interacting proteins relative to RMRP interacting proteins. Scatter plot of log2 transformed TMT ratios from two biological replicates is shown. Adjusted *P* value: two-tailed moderated *t*-test. **c**, Proteins enriched in SARS-CoV-2 RNA purifications (Supplementary Table 1). Left column: core SARS-CoV-2 RNA interactome (*P*<0.05). Left and right column: expanded SARS-CoV-2 RNA interactome. Significantly enriched proteins are highlighted in teal; SARS-CoV-2-encoded proteins are highlighted in pink. Adjusted *P* value: two-tailed moderated *t*-test.

To identify proteins that specifically interact with SARS-CoV-2 RNA, we compared the protein content of SARS-CoV-2 purifications to that of an unrelated control ribonucleoprotein complex of known composition. As the control, we selected the human RNA Component of Mitochondrial RNA Processing Endoribonuclease (RMRP) RNA and purified both SARS-CoV-2 and RMRP from HuH-7 cells 24 hours after infection. Covalent capture of purified proteins and sequencing of crosslinked RNA fragments confirmed that on average 90% of all mapped sequence reads originated from the SARS-CoV-2 genome in SARS-CoV-2 purifications (Supplementary Figure 1c). In RMRP purifications, on the other hand, more than 99% of crosslinked RNA fragments mapped to the human genome (Supplementary Figure 1c). We next analyzed the proteins purified in RAP-MS experiments by Western blot and detected the SARS-CoV-2 Nucleoprotein (N) only in SARS-CoV-2 purified samples (Supplementary Figure 1d). In contrast, the human RMRP component POP1 was discovered only in RMRP purifications (Supplementary Figure 1d). Together, these experiments demonstrate the specificity of the RAP approach for capturing the desired RNAs and their known direct protein binding partners.

### An unbiased atlas of direct SARS-CoV-2 RNA-protein interactions in human cells

Encouraged by the above results, we subjected proteins purified with both RMRP and SARS-CoV-2 RNAs to tandem mass tag (TMT) labeling and relative quantification by liquid chromatography coupled with tandem mass spectrometry (LC-MS/MS). In two biological replicate experiments, we identified 699 proteins, of which 583 were detected with two or more unique peptides (Supplementary Table 1). As shown in Figure 1b, we found five known RMRP components among the 10 most significantly enriched proteins in RMRP purifications (Supplementary Table 1). This is consistent with our previous results^13^, noting overall reduced RMRP expression levels in HuH-7 cells^20^.

Next, we quantified proteins enriched in SARS-CoV-2 purifications relative to RMRP purifications and found a total of 15 SARS-CoV-2 proteins, six of which were among the 20 most significantly enriched proteins (Figure 1b, c; Supplementary Table 1). In addition to five viral proteins translated from distinct open reading frames (ORFs), 10 of the 16 non-structural proteins (NSPs), which are derived from a precursor polyprotein^21^, were detected in RAP-MS experiments.

As expected, the SARS-CoV-2 N protein, which is thought to occupy the majority of the viral RNA, was one of the two most strongly enriched viral proteins, followed by several known viral RNA-binders, such as the endoribonuclease NSP15^22^, the RNA-dependent RNA polymerase (RdRP) NSP12^23^, the methyltransferase NSP16^24^, the single-stranded RNA-binding protein NSP9^25^, the capping factor NSP10^24^, the primase NSP8^23^, and the multifunctional protein NSP3^26^. In addition to NSPs, we also found ORF3A, which binds the 5’-end of the SARS-CoV-1 genomic RNA^27^, as well as the Spike protein (S), and the Membrane protein (M) among strongly enriched proteins.

While the M protein is known to interact with the RNA-binding protein N, a model for packaging of the coronavirus genomic RNA further suggests a possible direct RNA-binding function for M^28^. An RNA-binding activity of the S protein was not previously reported. While S covers the surface of the viral envelope, it has a transmembrane domain and an intracellular tail,^29^ making it conceivable that S may indeed contact viral RNA. Similarly, NSP3 and NSP6 are thought to be necessary for the formation of cytoplasmic double-membrane vesicles^30^, which contribute to viral replication or virion assembly and may contact RNA as part of these functions.

Finally, we also detected a protein corresponding to a newly annotated out-of-frame internal ORF (iORF) within the ORF of the N protein (N.iORF1)^31^. N.iORF1 shares 72% of its amino acids with ORF9b of SARS-CoV-1, making it likely that both proteins are homologous. N.iORF1 was moderately enriched in SARS-CoV-2 purifications (Figure 1b, c), suggesting that it may interact with viral RNA, which raises the possibility that at least two different RNA-binding proteins are translated from overlapping ORFs within the sequence of the viral N gene.

### Discovery of hundreds of SARS-CoV-2-bound human proteins

We next focused on the human proteins enriched in SARS-CoV-2 purifications. In total, we identified 276 proteins with a positive log2 fold-change in SARS-CoV-2 purified samples, relative to RMRP purified samples (Supplementary Table 1). Of these, 57 proteins were enriched with high statistical significance (*P* < 0.05, two-tailed moderated *t*-test), which we subsequently defined as the set of core SARS-CoV-2 interacting proteins (Figure 1c). Additionally, we also defined an expanded SARS-CoV-2 RNA interactome using a relaxed false discovery rate (FDR) of less than 20% (Figure 1c).

The expanded SARS-CoV-2 RNA interactome encompasses 104 human proteins and includes 13 SARS-CoV-2-encoded proteins. The vast majority of the human RNA interactome proteins (96 proteins, 92%) have previously been identified in high-throughput studies aimed at capturing proteins that crosslink to RNA^32^ (Supplementary Table 2). Comparing this expanded SARS-CoV-2 RNA interactome with the experimentally determined poly(A)-RNA interactome in HuH-7 cells^18^, revealed high overlap between both datasets (69 proteins, 66%) (Figure 2a; Supplementary Table 2). Next, we compared our direct SARS-CoV-2 RNA interactome with the group of proteins found to directly or indirectly associate with the RNA genomes of Dengue and Zika viruses^19^. Sixty-six proteins (63%) of the expanded SARS-CoV-2 RNA interactome were also found to associate with the Dengue and Zika virus RNAs, while 35 proteins (34%) were unique SARS-CoV-2 binders (Figure 2a). Since coronaviruses are known to form replication/transcription complexes (RTCs), we also compared the expanded SARS-CoV-2 RNA interactome to the protein content of murine coronavirus RTCs^33^ and found 64 shared proteins (Supplementary Table 2).

**Figure 2.**
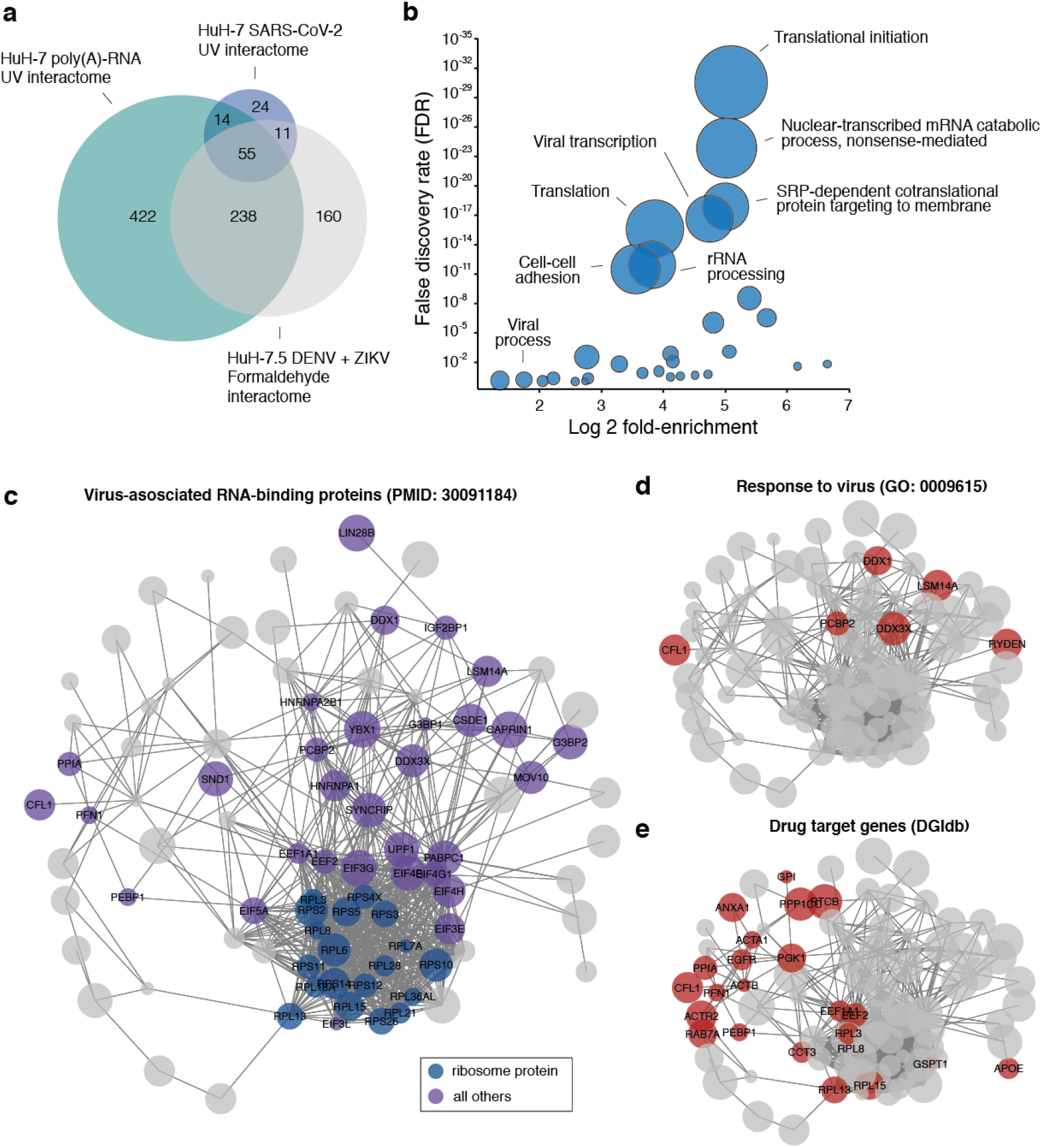
The SARS-CoV-2 RNA directly contacts regulators of RNA metabolism and host response pathways. **a**, Intersection of the expanded SARS-CoV-2 RNA interactome with the poly(A)-RNA interactome and the DENV/ZIKV interactome in HuH-7-derived cells. **b**, Gene ontology enrichment analysis of SARS-CoV-2 RNA interactome proteins. Circle sizes scale to the number of detected proteins. FDR calculation was performed using the Benjamini-Hochberg method. **c**, Protein-protein association network of SARS-CoV-2 enriched proteins. Published virus-associated proteins are highlighted. **d**, As in **c**, but proteins linked to the gene ontology term “response to virus” are highlighted. **e**, As in **c**, but proteins that overlap known drug target genes (DGIdb) are highlighted.

Finally, a recent study identified protein-protein interactors of recombinant SARS-CoV-2 proteins expressed in uninfected human cells^11^. Only 10 of the 334 human proteins that were found to interact with viral proteins in uninfected cells, also bound directly to viral RNA in infected cells (Supplementary Table 2). Beyond the well-documented impact of viral infection on host cell gene expression^5–7^, which may lead to remodeling of observed interactions, these results further highlight the importance of discriminating between protein-protein and RNA-protein interactions when dissecting the biology of SARS-CoV-2.

While our SARS-CoV-2 RNA interactome captured RNA-protein interactions not found in other RNA viruses or uninfected cells, many identified proteins were shared among these datasets, confirming the validity of our experimental approach. Further, the RNA-protein interactions only observed in SARS-CoV-2 RAP-MS experiments, may point towards unique aspects of SARS-CoV-2 biology.

### Biological functions of human SARS-CoV-2-bound proteins

To gain insights into the biological functions of SARS-CoV-2-bound proteins, we performed a hypergeometric gene ontology (GO) enrichment analysis^34^ on the expanded SARS-CoV-2 RNA interactome (see Methods). We observed strong enrichment for GO-terms linked to translational initiation (GO:0006413), nonsense-mediated decay (GO:0000184), SRP-dependent co-translational protein targeting to the membrane (GO:000661), and viral transcription (GO:0019083) (Figure 2b; Supplementary Table 3). Consistent with the enrichment of these GO-terms, the importance of mRNA translation at the endoplasmatic reticulum (ER) membrane is well-established for the replicative cycle of coronaviruses^35,36^. Further, nonsense-mediated RNA decay (NMD) was recently described as an intrinsic antiviral mechanism targeting coronavirus RNAs^37^.

In agreement with the crucial role of mRNA translation and its regulation, the expanded SARS-CoV-2 RNA interactome included 19 ribosomal proteins and 12 translation factors. Among translation factors, EIF4G1, which together with EIF4A and EIF4E, makes up the EIF4F complex, and EIF4B have been reported as targets of mammalian target of rapamycin (mTOR) signaling^38–40^. EIF4B plays a critical role in recruiting the 40S ribosomal subunit to mRNA, and both the PI3K/mTOR and mitogen-activated protein kinase (MAPK) pathways target EIF4B to control its phosphorylation status and activity^40^. PI3K/mTOR inhibition has further been demonstrated to suppress SARS-CoV-2 replication in human cells^8,10^.

In order to systematically examine the connectivity of the identified SARS-CoV-2 binding proteins and their potential relationship to virus-associated biological processes, we constructed a protein-protein association network of SARS-CoV-2 interacting proteins (Figure 2c; Supplementary Table 4; see Methods). Consistent with the crucial role of mRNA translation, ribosomal proteins and translation factors are prominently represented in this interaction network, along with previously published virus-associated RNA-binding proteins (Figure 2c) and proteins linked to the host response to viruses (Figure 2d).

We next integrated known drug-target interactions within this network and identified 23 SARS-CoV-2-bound human proteins that can be targeted with existing compounds, including PPIA, ANXA1, CFL1, and EGFR (Figure 2e). Notably, the core SARS-CoV-2 RNA interactome member ANXA1 plays an essential role as an effector of glucocorticoid-mediated regulation of the innate immune response. ANXA1 is a known target of dexamethasone, which according to newly emerging clinical trial data appears to show efficacy as a treatment for COVID-19^41^.

### The SARS-CoV-2 regulated proteome uncovers activated host response pathways

To complement our RNA-protein interactome and gain deeper insights into host response pathways activated upon SARS-CoV-2 infection, we globally measured protein abundance changes in infected cells. To this end, we performed triplicate MS experiments on SARS-CoV-2 infected and uninfected HuH-7 cells and identified 10,956 proteins with more than 2 unique peptides (Figure 3a; Supplementary Table 5). Among the detected proteins, 4,578 proteins were regulated (*P* < 0.05, moderated two-sample *t*-test) following 24 hours of SARS-CoV-2 infection, which is consistent with wide-spread proteome regulation upon SARS-CoV-2 infection^8^. As expected, proteome samples clustered according to their infection status in a principal component analysis (Supplementary Figure 2a). Among differentially expressed proteins, we detected 13 viral proteins (S, N, NSP3, N.iORF1, NSP2, NSP4, ORF3A, NSP1, ORF7A, M, ORF8, NSP8, and NSP10) and 56 proteins from our expanded SARS-CoV-2 RNA interactome (Figure 3a; Supplementary Table 5).

**Figure 3.**
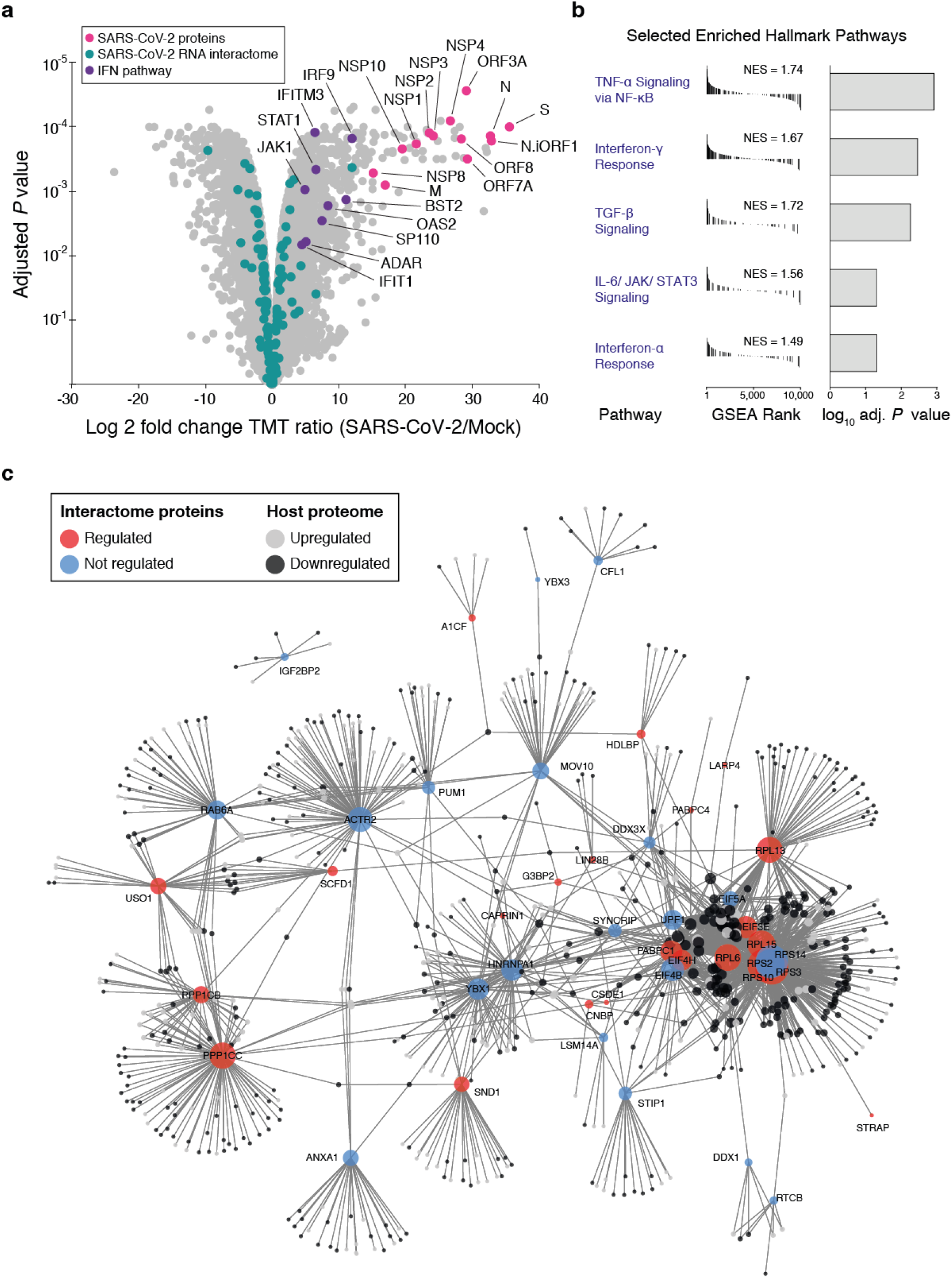
Connecting the SARS-CoV-2 RNA interactome to host cell perturbations upon infection. **a**, Volcano plot of proteome abundance measurements in SARS-CoV-2 infected and uninfected HuH-7 cells 24 hpi (*n*=3) (Supplementary Table 5). Adjusted *P* value: two-tailed moderated *t*-test. Pink: SARS-CoV-2-encoded proteins; Teal: human SARS-CoV-2-bound proteins; Purple: Interferon response-related proteins. **b**, Gene set enrichment analysis for global proteome abundance measurements shown in **a**. Selected gene sets are shown; full table displaying additional enriched gene sets is provided in Supplementary Figure 2b. **c**, Protein-protein association network of core SARS-CoV-2 RNA interactome proteins and their connections to differentially regulated proteins in SARS-CoV-2 infected cells, based on curated interactions in STRING v11^93^. Upregulated proteins are shown in light grey; downregulated proteins are shown in dark grey. Circle sizes scale to the number of connections of each interactome protein.

To identify cellular pathways modulated upon infection, we performed gene set enrichment analysis (GSEA) using our proteome abundance measurements (see Methods). Among the most significantly enriched hallmark gene sets were “TGFβ signaling” (adj. p = 0.0048), “TNFα signaling via NFκB” (adj. p = 0.0018), “Interferon (IFN)- γ response” (adj. p = 0.0048), “IL-6 JAK STAT3 signaling” (adj. p = 0.0054), and “IFN-α response” (adj. p = 0.0077; Figure 3b), suggesting a broad pro-inflammatory and antiviral response in infected cells. Further, we observed significant enrichment of the gene sets “GO Regulation of MAPK cascade” (adj. p = 0.00033), “GO Positive regulation of MAP Kinase activity” (adj. p = 0.011), and “GO Response to type I Interferon” (adj. p = 0.045; Supplementary Table 6). Newly emerging evidence indicates that these pathways are indeed highly relevant in the context of SARS-CoV-2 infections^5,7–10,42^. Notably, inhibition of growth factor signaling through the MAPK pathway, which responds to and controls the production of pro-inflammatory cytokines, including TNFα and IL-6, was recently shown to modulate SARS-CoV-2 replication in infected cells^8–10^.

In agreement with recent transcriptome studies^5–7^, our proteome data suggest activation of interferon signaling upon SARS-CoV-2 infection. Among interferon-related genes, we observed significant upregulation of several major components of IFN signaling pathways, including STAT1 and IRF9, which together with STAT2 make up the ISGF3 complex, their upstream components TYK2 and JAK1, as well as their downstream targets IFIT1, IFIT3, IFITM3, OAS2, and ISG15 (Figure 3a). Other strongly upregulated IFN-related genes include BST2, SP110, UBE2L6, ADAR, TGIF1, BRD3, and IFI30 (Supplementary Table 5). Notably, many SARS-CoV-2 RNA interactome members are directly or indirectly linked to the IFN response. These include PUM1^43,44^, YBX1^44,45^, SYNCRIP^46^, G3BP1^47–50^, G3BP2^47,48^, EIF4B^51^, MOV10^52–54^, CAPRIN1^48^, DDX3X^55–58^, LSM14A^59,60^, RYDEN^61,62^, STRAP^63^, ANXA1^64,65^, DDX1^66–68^, PCBP2^69,70^, HNRNPA2B1^71^, and YWHAZ^72^. In conclusion, our global protein abundance measurements verify the induction of an appropriate host response in SARS-CoV-2 infected HuH-7 cells and further support an essential role for IFN and MAPK signaling in SARS-CoV-2 infections.

### Connectivity map of SARS-CoV-2 RNA binders and regulated host cell proteins highlights regulators of emerging SARS-CoV-2 biology

To examine the connectivity between SARS-CoV-2 RNA interactome proteins and proteins that are differentially regulated in host cells upon viral infection, we used curated protein-protein interaction data to build a network that visualizes interactions between SARS-CoV-2 RNA binders and regulated host proteins (Figure 3c and Supplementary Figure 2c; Supplementary Table 7; see Methods). In addition to a cluster of ribosomal proteins and translation factors, we observed many connections between canonical RNA-binding proteins linked to viral infections (including MOV10, YBX1, HNRNPA1, PUM1, DDX3X, HDLBP, SND1, and CFL1)^73^ and regulated host factors in this network. Notably, we also observed non-classical RNA-binding proteins such as ANXA1 or ACTR2 among highly-connected proteins in this network. ACTR2 has been identified as a host factor for respiratory syncytial virus (RSV) infection and is involved in cytoskeleton reorganization and filipodia formation, which is thought to promote virus spreading^74,75^. This observation is consistent with recent work that demonstrated that SARS-CoV-2 infection induced a dramatic increase in filopodia, and viral particles localized to and emerged from these protrusions^9^.

Taken together, this network analysis suggests extensive connectivity between SARS-CoV-2-binders and dynamically regulated host cell proteins, which connect interactome proteins to emerging SARS-CoV-2 biology and may provide a map for identifying regulatory hubs in SARS-CoV-2 infections.

### The transcriptional regulator CNBP directly engages the SARS-CoV-2 RNA

We next focused our analysis on SARS-CoV-2 RNA interactome members that were strongly upregulated in our proteome dataset following viral infection. Among the 104 proteins in the expanded SARS-CoV-2 RNA interactome, 22 human proteins were significantly induced upon infection. In this group, PPIA, RAB6D, MSI2, SCFD1, and CNBP were the most strongly upregulated candidates (log2 fold-change > 2.5) and all of these proteins also showed robust enrichment in RAP-MS experiments (log2 fold-change > 1, *P* <0.1). Notably, PPIA was previously shown to be essential for SARS-CoV-1 replication^76,77^, suggesting that the group of virally induced human SARS-CoV-2 interactors may constitute important candidates for a functional investigation.

To further dissect these candidates and validate our RAP-MS results, we first focused on CNBP, which represents the most significantly enriched protein in SARS-CoV-2 purifications and was also induced upon infection. CNBP translocates to the nucleus upon immune stimulation and associates with an immune stimulatory DNA oligonucleotide, suggesting that it may be involved in foreign nucleic acid-sensing pathways^78^. Two recent studies showed that CNBP activates the innate immune response and induces the expression of several pro-inflammatory cytokines, including Il-6 and Cxcl10 *in vivo*^78,79^. While our proteomics experiments did not yield peptides for these two CNBP-regulated cytokines, recent transcriptome studies demonstrated that IL-6 and CXCL10 were indeed induced in SARS-CoV-2 infected human cells^5–7^. In light of these results, we speculated that in addition to its previously described role in foreign DNA sensing, CNBP might recognize the SARS-CoV-2 RNA through direct binding, which may be linked to its role in transcriptionally activating specific innate immune genes.

To corroborate the physical engagement of the SARS-CoV-2 RNA by CNBP in infected human cells, we combined RNA antisense purification (RAP) with enhanced crosslinking and immunoprecipitation (eCLIP)^80^. Briefly, following the purification of SARS-CoV-2-bound proteins from infected HuH-7 cells using RAP, we incubated eluted proteins with an antibody recognizing CNBP, or an IgG antibody as negative control. Immunoprecipitations (IPs) were subjected to denaturing purification using SDS-PAGE, and size-matched RNA populations were extracted from CNBP and IgG samples and converted into cDNA libraries. As shown in Figure 4a, we observed strong binding of CNBP to the viral RNA genome, while only minimal signal was observed in size-matched IgG libraries, suggesting specific binding of CNBP to the SARS-CoV-2 RNA. These data provide strong evidence for a direct interaction between CNBP and the SARS-CoV-2 RNA in infected human cells and validate that RAP-MS indeed recovers direct SARS-CoV-2 binders.

**Figure 4.**
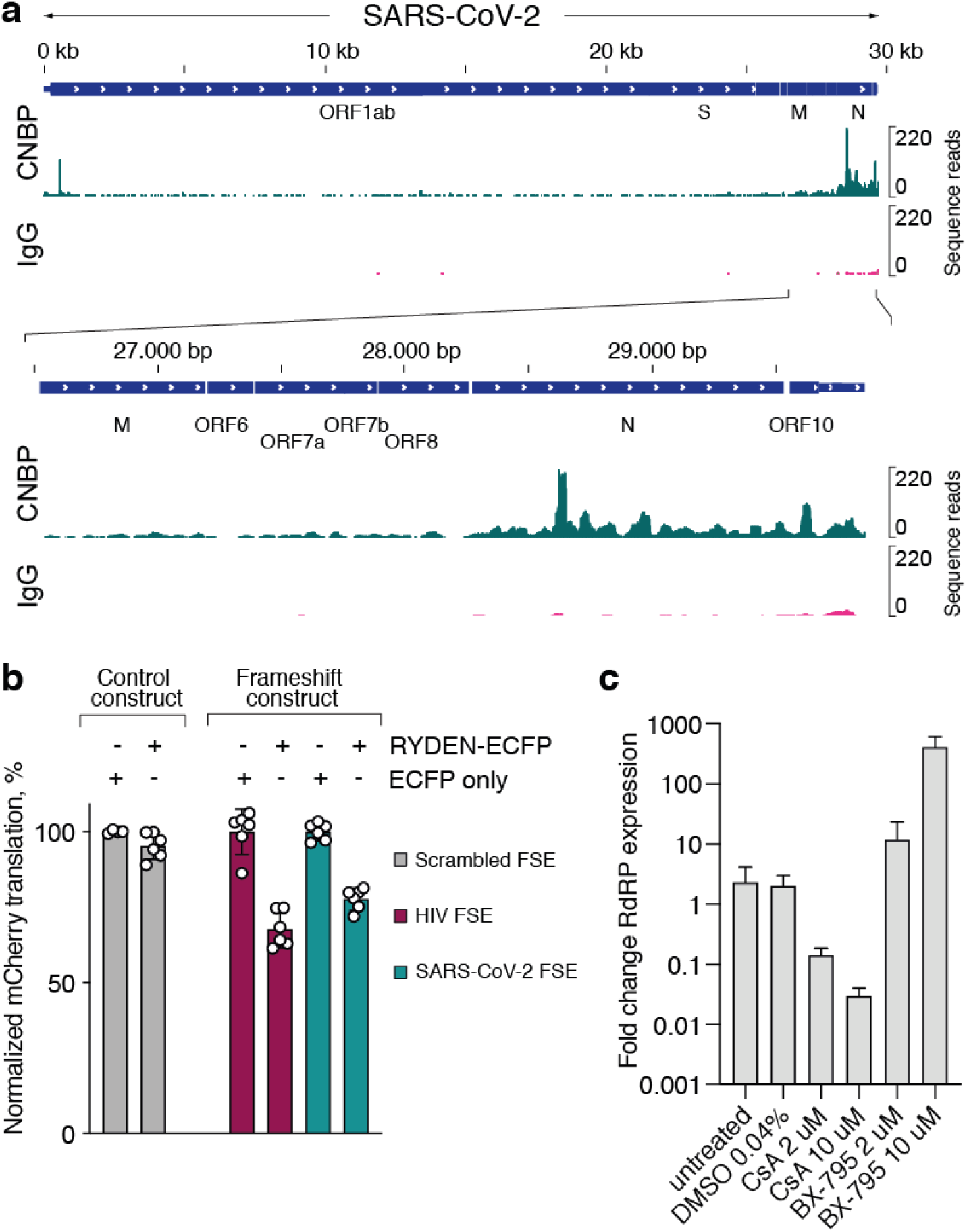
RAP-MS uncovers direct and physiologically relevant RNA-protein interactions. **a**, CNBP eCLIP-seq data plotted across the SARS-CoV-2 genome. Control libraries using an unspecific IgG antibody are sown below CNBP alignments. Zoom-in to CNBP-bound region at the 3’-end of the SARS-CoV-2 genome is sown in the lower panel. **b**, Quantification of ribosomal frameshifting efficiency using a dual-fluorescence translation reporter (Supplementary Figure 3a), containing either a scrambled FSE, the HIV FSE or the SARS-CoV-2 FSE. Normalized to cells transfected with ECFP. Values are mean ± standard deviation (*n*=6, except for control RNA *n*=4). **c**, RT-qPCR measurements of genomic intracellular SARS-CoV-2 RNA (RdRP gene) at 72 hpi in HuH-7 cells treated with or without the indicated inhibitors for 2 h prior to SARS-CoV-2 infection. Values are mean ± SEM (*n*=3). FSE: frameshift element.

### RYDEN suppresses ribosomal frameshifting during translation of the SARS-CoV-2 RNA

Following CNBP, the second most significantly enriched human protein in our SARS-CoV-2 RNA interactome was LARP4, a known regulator of mRNA stability and translation^81^. LARP4 and LARP1 interact with PABPC1^81^, and both LARP1 and PABPC1 have been proposed to reside in the same ribonucleoprotein (RNP) complex with RYDEN^61^, all of which were enriched in RAP-MS experiments. RYDEN, in particular, attracted our attention. Similar to CNBP, RYDEN was among the most significantly enriched proteins in SARS-CoV-2 purifications and was also significantly upregulated upon infection. In addition to being an IFN-induced gene, RYDEN was previously shown to suppress virus production in dengue virus (DENV) infected cells^61^, and has further been demonstrated to inhibit programmed –1 ribosomal frameshifting (−1FS) in human immunodeficiency virus (HIV) infections^62^. To execute this function, RYDEN requires the eukaryotic peptide chain release factor GSPT1^62^, which was detected in our expanded RNA interactome as well (Figure 1b, c).

To dissect if RYDEN can modulate the frequency of −1FS in SARS-CoV-2, we generated a dual-color fluorescence reporter system that allowed us to quantify frameshifting efficiency in the presence or absence of RYDEN. In brief, cells transfected with this reporter express a single fluorescent protein (EGFP) when the 0 reading frame is translated (Supplementary Figure 3a). Expression of a second fluorescent protein (mCherry) downstream of EGFP is dependent on −1FS, which prevents translation of an in-frame stop codon. Thus, the ratio between mCherry and EGFP directly correlates to −1FS efficiency. As a normalization control, we used a construct lacking a stop codon in the 0 reading frame, leading to the expression of EGFP and mCherry in equal ratios (Supplementary Figure 3a). Similar to other coronaviruses, SARS-CoV-2 frameshifting is very efficient, reaching 33% in human HEK293 cells. Using a reporter containing the HIV frameshift element (FSE) as a positive control, we first confirmed that overexpression of a RYDEN-ECFP fusion protein suppressed −1FS when compared to ECFP expression alone (Figure 4b). Strikingly, overexpression of RYDEN also led to a significant reduction of ribosomal frameshifting efficiency, when a reporter containing the SARS-CoV-2 FSE was used (Figure 4b). Together, these results show that RYDEN (i) is induced upon SARS-CoV-2 infection, (ii) associates with the SARS-CoV-2 RNA in infected cells, and (ii) inhibits programmed ribosome frameshifting of the SARS-CoV-2 FSE.

### Inhibiting components of the human SARS-CoV-2 RNA-protein interactome modulates viral replication

Finally, we set out to test if targeting components of our SARS-CoV-2 RNA interactome and its associated pathways with known inhibitors proved effective in manipulating intracellular viral replication. To this end, we treated HuH-7 cells with the PPIA inhibitor Cyclosporin A (CsA) and observed a dose-dependent reduction in the expression of intracellular viral RNA (Figure 4c). The treatment with CsA had no apparent effect on cell viability (Supplementary Figure 3b). PPIA is involved in the folding of proteins and has a well-documented impact on different viruses^82^. Yet, its direct interaction with the SARS-CoV-2 RNA genome has not been demonstrated and expands these previously described functions. Notably, PPIA represents the most strongly induced interactome member in SARS-CoV-2 infected cells.

Given the enrichment of various proteins linked to the IFN response in our SARS-CoV-2 RNA interactome, we speculated that inhibiting a central regulator of antiviral RNA sensing might make HuH-7 cells more permissive to SARS-CoV-2 replication. We noticed that several SARS-CoV-2 interacting proteins engage in well-documented physical interactions with TANK-binding kinase (TBK1). Of particular note, DDX3X is a TBK1 substrate and direct binding partner that synergizes with TBK1 in regulating IFN production^57^. Similarly, ANXA1 regulates type I IFN signaling through its physical association with TBK1^65^, while STRAP interacts with TBK1 and IRF3 to promote IFN-β production^63^. Indeed, treatment of HuH-7 cells with the TBK1 inhibitor BX-795 increased intracellular virus replication by a factor of several hundred-fold, suggesting a diminished host-response. Inhibitor treatment only had a small effect on cell viability at the tested concentrations (Supplementary Figure 3b). These results demonstrate that proteins and pathways linked to the direct SARS-CoV-2 RNA interactome identified in this work, represent viable targets for inhibiting coronavirus replication or modulating the host cell response to viral infections. The SARS-CoV-2 RNA interactome provides valuable starting points for mechanistic studies of viral infection and may further be explored for evaluating treatment options for COVID-19.

## DISCUSSION

Decoding the RNA-protein interactomes of pathogenic RNA viruses has been a long-standing challenge. In the face of the current COVID-19 pandemic, the task of addressing this challenge has gained considerable attention and urgency. In this study, we provide detailed molecular insights into the identity of host factors and machinery that directly engage the SARS-CoV-2 RNA during infection of human cells. Using RAP-MS, we have obtained an unbiased and quantitative picture of direct contacts between the host cell proteome and viral RNA. Combining this global RNA-protein interaction atlas with proteome-wide measurements of protein abundance changes that occur upon SARS-CoV-2 infection, allowed us to chart the RNA-protein interaction landscape of the SARS-CoV-2 RNA in infected human cells, and illuminate how these interactions intersect with host cell perturbations and host defense pathways.

In addition to the induction of an IFN response, we provide evidence for the activation of MAPK and TNFα signaling pathways upon SARS-CoV-2 infection, which adds to growing evidence that these pathways are regulated upon SARS-CoV-2 infection and may be exploited to manipulate viral replication^8–10^. In the context of EGF/MAPK signaling, it is intriguing that we detected EGFR together with several RAS-related proteins, including RAB6D, RAB6A and RAB7A in our expanded RNA interactome. RAS-related proteins were also identified as protein-protein interaction partners of viral proteins in human cells, including RAB7A, which we find to bind to the SARS-CoV-2 RNA in this study. Together, these data may point at physical interactions between SARS-CoV-2 components and cellular pathways that are perturbed upon viral infection.

Beyond characterizing the SARS-CoV-2 RNA interactome, our work uncovered a subset of SARS-CoV-2 binders that are induced upon viral infection. Importantly, this group of proteins includes a previously identified host dependency factor essential for coronavirus replication (PPIA), suggesting that our approach indeed recovers relevant regulators of coronavirus biology. We characterized two of these proteins in greater detail. First, we demonstrated that CNBP directly interacts with the SARS-CoV-2 RNA, which expands its previously known role in the context of foreign DNA sensing. It is tempting to speculate that recognition of the SARS-CoV-2 RNA by CNBP might contribute to establishing a specific cytokine response in infected cells. Intriguingly, the known CNBP-dependent cytokine expression signature^78,79^ bears similarity to gene expression changes in SARS-CoV-2 infected cells^5–7^, which warrants future investigations. In addition to CNBP, we provide evidence that the interferon-induced protein RYDEN interacts with SARS-CoV-2 RNA and suppresses programmed ribosome frameshifting in a SARS-CoV-2 translational reporter. Together, these insights illuminate how host cell proteins directly engage foreign RNA molecules during infection and contribute to different host defense strategies.

Various additional host cell proteins can act as effector molecules in such defense mechanisms. RNA editing by host deaminases for instance is an innate restriction process to counter virus infection. Recent work revealed evidence for host-dependent RNA editing in SARS-CoV-2^83^. Consistent with these reports we find the APOBEC1 complementation factor (A1CF) among the most significantly enriched SARS-CoV-2 interactors. Further, the functionally related adenosine deaminase acting on RNA (ADAR) is an interferon-induced gene that was strongly upregulated in response to SARS-CoV-2 infection in our proteome analysis.

Another notable group of human SARS-CoV-2 interactors are proteins associated with various vesicle trafficking functions, including SCFD1, USO1, RAB1A, RAB6D, RAB6A, and GDI2. The significant enrichment of these proteins in the SARS-CoV-2 RNA interactome is particularly interesting, given the crucial roles of membranes and vesicles for the formation of RTCs and virion assembly. We further detect two viral proteins involved in double-membrane vesicle formation (NSP3 and NSP6) in our RNA interactome. This highlights the possibility that an RNA-binding activity of various membrane or vesicle-associated proteins may promote packaging of the viral RNA into newly assembling virions or membrane vesicles.

Our characterization of the SARS-CoV-2 RNA interactome is a valuable resource that provides an RNA-centric perspective on SARS-CoV-2 infections and will help to decode both viral RNA metabolism as well as host defense mechanisms. As demonstrated in our study, the identified interactions recover known coronavirus biology and uncover opportunities for targeting proteins or pathways linked to the SARS-CoV-2 RNA interactome in order to control viral replication. We believe that our findings and strategies provide a general roadmap for dissecting the biology of RNA viruses and the interactions between hosts and pathogens at the molecular level.

## MATERIALS AND METHODS

### Tissue culture

We maintained HuH-7 cells in DMEM media (Thermo Fisher Scientific) supplemented with 10% heat-inactivated FBS (Thermo Fisher Scientific), and 100 units/ml streptomycin and 100 mg/ml penicillin. Cells were grown at 37°C and 5% CO_2_ atmosphere.

### Virus production

We used a previously described patient-derived SARS-CoV-2 isolate^84^ that was propagated on HuH-7 cells and characterized by RNA sequencing. Viral loads were frequently determined by RT-qPCR. In brief: viral RNA was extracted with a MagNA Pure 24 system (Roche, Germany) and quantified with the LightMix Assay SARS-CoV-2 RdRP RTqPCR assay kit (TIB MOLBIOL, Germany) and the RNA Process Control kit (Roche). Viral titers were determined by immunofluorescence.

### RAP-MS

RAP-MS was carried out as previously described^13^ with the following modifications: To capture endogenous SARS-CoV-2 transcripts, we designed and synthesized 5’ biotinylated 90-mer DNA oligonucleotides (Integrated DNA Technologies) antisense to the complementary target RNA sequence. We used 67 probes such that one probe binding site occurs roughly every 400 bases in the ~30 kb SARS-CoV-2 genome and excluded regions that matched to human transcripts or genomic regions as previously described^85,86^. For SARS-CoV-2 and RMRP antisense purifications, we grew ten 10 cm tissue culture plates of HuH-7 cells per replicate. We prepared two replicates of each of the SARS-CoV-2 and RMRP samples and included two non-crosslinked control samples that were used for RMRP purifications. SARS-CoV-2 infection was carried out with an MOI of 10 for 24 hours. Cells were washed once with PBS and then crosslinked on ice using 0.8 J/cm^2^ of 254 nm UV light in a GS Gene Linker (Bio-Rad). Cells were then lysed on the tissue culture plate by adding 1 ml of RAP lysis buffer (10 mM Tris pH 7.5, 500 mM LiCl, 0.5% dodecyl maltoside (DDM), 0.2% sodium dodecyl sulphate (SDS), 0.1% sodium deoxycholate, EDTA-free Protease Inhibitor Cocktail (Thermo Fisher Scientific) and Murine RNase Inhibitor (New England Biolabs)). Lysates were then collected and flash-frozen in liquid nitrogen for storage at −80°C. All subsequent steps were carried out at previously described^13^.

### RAP-MS protein digestion and TMT labeling

RAP-captured proteins were resuspended in 40 μL of 8 M urea in 50 mM Tris-HCl, followed by reduction with 4 mM DTT for 30 minutes at room temperature and alkylation with 10 mM IAA for 45 minutes at room temperature in the dark. All six samples were then digested with 0.1 μg Lys-C for 2 hours, followed by a reduction of the urea concentration to <2 M and continued digestion with 0.5 μg trypsin overnight. Reactions were quenched with formic acid at a final concentration of 5% and then desalted by reverse phase C18 stage tips as described previously^87^ and dried down. Peptides were then resuspended in 50 μL of 50 mM HEPES buffer and isobarically labeled using 400 μg of Tandem Mass Tag 6-plex (TMT6) isobaric labeling reagent (Thermo Fisher Scientific). The labeling reactions were then quenched with 4 μL of 5% hydroxylamine, samples were mixed together, and dried. The sample was fractionated by SCX stage tip strategy using three pH cuts at 5.15, 8.25, and 10.3 as described previously^87^.

### Proteome analyses of SARS-CoV-2 infected cells

For proteome measurements, we expanded HuH-7 cells to two 10cm tissue culture plates per replicate. Cells were infected with an MOI of 10 and incubated for 24 hours before harvesting. Three process replicates of infected and non-infected cell line samples were generated. Cells were lysed in 8 M Urea, 75 mM NaCl, 50 mM Tris pH 8.0, 1mM EDTA, 2 μg/ml Aprotinin, 10 μg/ml Leupeptin, 1 mM PMSF, 10 mM NaF, PIC2 (Sigma Aldrich), PIC3 (Sigma Aldrich) and 10mM Sodium Butyrate. Benzonase was added to digest nucleic acids and DNA was sheared using a probe sonicator (10% amplitude, 0.7s on, 2.3s off, 6 min 15s total). Cell debris was removed by centrifugation and lysates were flash-frozen for storage. All samples were prepared for MS analysis using an optimized workflow as described previously^88^. Briefly, lysed samples were reduced, alkylated, and digested by LysC for 2 hours, followed by overnight digestion with trypsin. Digestions were quenched with formic acid and all peptide samples were desalted using reverse phase C18 SepPak cartridges. Samples were then quantified by the Pierce BCA Protein Assay and measured into 500 μg aliquots for isobaric labeling. Peptides were isobarically labeled with TMT6 following the reduced TMT protocol^89^. After confirming 98% or greater label incorporation, the samples were mixed together and desalted. Sample was then fractionated by offline high pH reversed phase chromatography and concatenated into 24 fractions for the analysis by on-line LC-MS/MS^88^.

### LC-MS/MS analysis (RAP-MS and proteome)

All the samples were analyzed either on an Orbitrap Exploris™ 480 (RAP-MS fractions) or Q Exactive ™ Plus (proteome fractions) mass spectrometer coupled with Easy-nLC 1200 ultra-high pressure liquid chromatography (UPLC) system (Thermo Fisher) with solvent A of 0.1% formic acid (FA)/3% acetonitrile and solvent B of 0.1% FA/90% acetonitrile. One μg of each of the proteome fractions and half of each of the RAP-MS fractions were injected on a 75μm ID Picofrit column packed in-house to approximately 28 cm length with Reprosil-Pur C18-AQ 1.9 μn beads (Dr. Maisch GmbH). Samples were separated at 200 nL/min flow rate with a gradient of 2-6% solvent B for 1min, 6-30% B in 84 min, 30-60% B in 9 min, 60-90% B in 1 min, followed by a hold at 90% B for 5 min. Both mass spectrometers were operated in data dependent acquisition mode. On Exploris 480 MS1 scan (r = 50,000) was followed by MS2 scans (r = 15,000) for top 20 most abundant ions using normalized automatic gain control (AGC) of 100% for MS1 and 200% for MS2, MS2 maximum inject time of 150ms, normalized collision energy of 34 and fit filter of 50%. On Q Exative Plus MS parameters were set as follows: MS1 r = 70,000; MS2 r = 17,500; MS1 AGC target of 3e6, MS2 for 12 most abundant ions using AGC target of 5e4 and maximum inject time of 120ms, and normalized collision energy of 29.

### Quantification and identification of peptides and proteins (RAP-MS and proteome)

MS/MS spectra were searched on the Spectrum Mill MS Proteomics Workbench against a RefSeq-based sequence database containing 41,457 proteins mapped to the human reference genome (hg38) obtained via the UCSC Table Browser (https://genome.ucsc.edu/cgi-bin/hgTables) on June 29, 2018, with the addition of 13 proteins encoded in the human mitochondrial genome, 264 common laboratory contaminant proteins, 553 human non-canonical small open reading frames, 28 SARS-CoV-2 proteins obtained from RefSeq derived from the original Wuhan-Hu-1 China isolate NC_045512.2^90^, and 23 novel unannotated SARS-CoV-2 ORFs whose translation is supported by ribosome profiling^31^, yielding a total of 42,337 proteins. Amongst the 28 annotated SARS-CoV-2 proteins we opted to omit the full length ORF 1ab, to simplify peptide-to-protein assignment, and instead represented ORF1a and ORF1ab as the mature 16 individual non-structural proteins that result from proteolytic processing of the 1a and 1ab polyprotein. We further added the D614G variant of the SARS-CoV-2 Spike protein that is commonly observed in European and American virus isolates. Spectrum Mill search parameters included: instrument setting of ESI-QEXACTIVE-HCD-v4-35-20, parent and fragment mass tolerance of 20ppm, trypsin allow P enzyme setting and up to 4 missed cleavages. Carbamidomethylation and TMT labeling at Lysine (with and without labeling at N-terminus) were set as fixed modifications, while variable modifications included acetylation of protein N-termini, oxidized methionine, deamidation of asparagine, and pyro-glutamic acid at peptide N-terminal glutamine. Peptide spectrum match score thresholding was optimized to achieve a target-decoy false discovery rate (FDR) of 1.2% for validation of spectra. Peptide level auto-validation was followed by protein polishing with FDR of 0% at protein level and minimum score of 13.

The Spectrum Mill generated proteome level export from the RAP-MS and proteome datasets filtered for human proteins identified by two or more distinct peptides, SARS-CoV-2 proteins and unannotated virus ORFs were used for further statistical analyses. Five of the SARS-CoV-2 non-structural proteins (NSP6, NSP15, NSP16, NSP9, and NSP1) identified by a single, highly scoring distinct peptide were kept in the dataset. Keratins were excluded from RAP-MS data. Protein quantification was achieved by taking the ratio of TMT reporter ion for each sample/channel over the median of all 6 channels. Moderated two-sample *t*-test was applied to compare SARS-CoV-2 and RMRP samples after mean normalization and SARS-CoV-2 infected and non-infected samples after median-MAD normalization of RAP-MS and proteome datasets, respectively. Benjamini-Hochberg corrected p-value threshold of 0.05 was used to asses significantly regulated proteins in each of the datasets.

### Western blot of RAP-MS samples

Following the described RAP-MS procedure, we saved 5% of benzonase eluted proteins for western blot analysis. We added NuPAGE LDS Sample Buffer (Thermo Fisher Scientifc) and incubated samples for 3 min incubation at 95°C. Proteins were resolved by SDS-PAGE using NuPAGE 4–12% Bis-Tris-HCl Gels (Thermo Fisher Scientific) at 200 V for 1 h, followed by transfer to a nitrocellulose membrane using the iBlot Dry Blotting System (Thermo Fisher Scientific). Western blots were performed using the iBind Western System (Thermo Fisher Scientific). For protein detection, we used the following primary antibodies: Nucleoprotein – Abcam #ab272852, POP1 – Proteintech #12029-1-AP. We used the following secondary antibodies: IRDye 800CW Goat anti-Rabbit IgG (H + L) (LI-COR), IRDye 800CW Donkey anti-Goat IgG (H + L) (LI-COR). For visualization of bands, we used the Odyssey Clx infrared imager system (LI-COR).

### Covalent protein capture and sequencing of crosslinked RNA

To capture RNA sequences covalently crosslinked to proteins purified with RAP-MS, we carried out RNA antisense purifications as described above. Following our pilot RAP-MS experiment (see Supplementary Figure 1a, b), SARS-CoV-2 bound proteins were eluted from streptavidin beads by heat fragmentation of RNA (3 minutes at 91°C in 100 mM HEPES pH 7.5, 5 mM MgCl_2_, 100 mM KCl, 0.02% Triton X-100). For subsequent RAP-MS experiments we replaced heat fragmentation with RNase H digestion, using 7.5 μl RNase H (NEB), 2 μl TURBO DNase (Thermo Fisher Scientific) in 55.5 μl RNase H buffer (100 mM HEPES pH 7.5, 75 mM NaCl, 3 mM MgCl_2_, 0.125% NLS, 0.025% sodium deoxycholate, 2.5 mM TCEP) and incubating 30 minutes at 37°C. Following elution of proteins, supernatants were transferred into new tubes and beads were washed once with RNase H buffer. Wash fractions were pooled with eluates and stored on ice. The next steps were previously described in similar form by Quinodoz et al^91^. We separated 100μl of NHS magnetic beads (Thermo Fischer) on a magnet and discarded the supernatant. We then washed with 1 ml 1 mM ice cold HCl, followed by a quick rinse in 1 ml ice-cold PBS. After removing PBS, we immediately added the stored eluates to the prepared beads. Binding was carried out overnight at 4°C on a rotating wheel. The next day, we quenched NHS beads by adding 1 ml of 0.5 M Tris pH 8.0 and incubating for 1h at 4°C. We then washed beads four times in 1 ml of Modified RLT buffer (RLT buffer supplied by Qiagen with added 10 mM Tris pH 7.5, 1 mM EDTA, 1 mM EGTA, 0.2% NLS, 0.1% Triton-X, and 0.1% NP-40). Next, we washed beads two more times in 1 ml 1X PBS, 5 mM EDTA, 5 mM EGTA, 5 mM DTT, 0.2% Triton-X, 0.2% NP-40, 0.2% sodium deoxycholate and incubated each washing step 5 minutes at 55°C. These heated washing steps were followed by two additional washed in the same buffer at room temperature. Subsequently, beads were rinsed on the magnet in 1x FastAP buffer (10 mM Tris pH 7.5, 5 mM MgCl_2_, 100 mM KCl, 0.02% Triton X-100). Next, end repair was carried out by resuspending beads in 50 μl FastAP mix (39 μl H_2_O, 5 μl 10x FastAP buffer (Thermo Fisher Scientific), 1 μl Murine RNase Inhibitor (NEB), 5 μl FastAP enzyme (Thermo Fisher Scientific) and incubating 20 minutes at 37°C. In the meantime, we prepared 150 μl of T4 PNK mix (120 μl H_2_O, 20 μl 10x T4 PNK buffer (NEB), 1 μl Murine RNase Inhibitor (NEB), 7μl T4 PNK (NEB), 1 μl TURBO DNase (Thermo Fisher Scientific)), which was added to the FastAP reaction and incubated for another 20 minutes at 37°C. Following end repair, we washed beads once in Modified RLT buffer, and two times in Detergent Wash Buffer (20 mM Tris pH 7.5, 50 mM NaCl, 0.2% Triton-X, 0.2% NP-40, 0.2% sodium deoxycholate). We then rinsed beads on the magnet twice with 1X T4 RNA ligase buffer (50 mM Tris-HCl pH 7.5, 10 mM MgCl_2_), before resuspending the beads in 25 μl RNA ligation mix (9 μl H_2_O, 3 μl 10x T4 RNA ligase buffer (NEB), 0.3 μl 0.1M ATP, 0.8 μl DMSO, 0.4 μl Murine RNase Inhibitor (NEB), 9 μl PEG 8000, 2.5 μl T4 RNA ligase I High Concentration (NEB). Next, we added 5 μl 20 nM RiL19 (/5phos/rArGrArUrCrGrGrArArGrArGrCrGrUrCrGrUrG/3SpC3/; IDT) and incubated samples for 75 minutes at 23°C. Following 3’-ligation, we washed the beads once in 1ml Modified RLT Buffer, followed by two washes in Detergent Wash Buffer. Next, we resuspended beads in 250 μl Proteinase K mix containing 200 μl NLS RNA Elution buffer (20 mM Tris pH 8.0, 10 mM EDTA, 2% NLS, 2.5 mM TCEP), 12.5 μl 5 M NaCl, 1 μl 500 mM TCEP, 12.5 μl Proteinase K (NEB), and 24 μl H_2_O and incubated samples for 1.5 hours at 55°C. Following Proteinase K digestion, we separated beads on a magnet, transferred the supernatant into a new tube and extracted RNA using Phenol/Chloroform extraction. All subsequent manipulation steps were carried out as described in the eCLIP library preparation protocol^80^, starting with the reverse transcription of recovered RNA fragments.

### eCLIP

To facilitate eCLIP experiments in SARS-CoV-2 infected HuH-7 cells, we applied the previously described RAP-MS approach to purify all SARS-CoV-2-bound proteins prior to immunoprecipitating individual proteins from this pool. Similar strategies were previously applied to facilitate eCLIP experiments^92^.

Following RAP-MS, SARS-CoV-2-bound proteins were eluted by adding 7.5 μl RNase H (NEB), 2 μl TURBO DNase (Thermo Fisher Scientific) to 55.5 μl RNase H buffer (100 mM HEPES pH 7.5, 75 mM NaCl, 3 mM MgCl_2_, 0.125% NLS, 0.025% sodium deoxycholate, 2.5 mM TCEP) and incubating 30 minutes at 37°C. Subsequently, we adjusted the volume of eluates to 250 μl with H_2_O and added an equal volume of 2x CLIP lysis buffer (100 mM Tris-HCl pH 7.4, 300 mM NaCl, 2 mM EDTA, 2% (v/v) NP40, 1% sodium deoxycholate, 0.5 mM DTT). In the meantime, we coupled 100 μl Protein G Dynabeads (Thermo Fisher Scientific) with 10 μg CNBP antibody (Proteintech, 14717-1-AP) or 10 μg IgG antibody (Cell Signaling Technology, 2729) by incubating antibody-bead suspensions 45 minutes at room temperature. We removed unbound antibody and added the stored eluates to antibody-coupled beads and incubated the beads for 4 hours at 4 °C. Following incubation, immunoprecipitates were washed two times in 1 ml CLIP lysis buffer, two times in IP wash buffer (50 mM Tris-HCl pH 7.4, 300 mM NaCl, 1 mM EDTA, 1% (v/v) NP40, 0.5% sodium deoxycholate, 0.25 mM DTT), followed by two washes in 50 mM Tris-HCl pH 7.4, 1 mM EDTA, 0.5% (v/v) NP40. All subsequent manipulation steps were carried out according to the eCLIP protocol^80^.

### RNA extraction and RT-qPCR

At 72 h post-infection, cells were lysed in 3 volumes RLT buffer (Qiagen) and RNA was extracted using Dynabeads MyOne Silane (Life technologies). RNA was bound to the beads by adding 4.5 volumes isopropanol, washed twice with 70% EtOH, DNase treated (TURBO DNase, Thermo Fisher Scientific), washed again with 70% Et-OH and eluted in H20. RNA was reversely transcribed into cDNA (SmartScribe RT, Takara Bio) according to the manufacturer’s instructions. Viral RNA was quantified by qPCR using PowerUp SYBR Green master mix (Thermo Fisher Scientific) and primers specific to the SARS-CoV-2 RdRP gene (fwd: GTGARATGGTCATGTGTGGCGG, rev: CARATGTTAAASACACTATTAGCATA) and 18S rRNA (fwd: ATGGCCGTTCTTAGTTGGTG, rev: GAACGCCACTTGTCCCTC TA). We calculated differences in RNA expression using the ΔΔCT method versus 18S. To achieve power to detect small effects in gene expression, we performed 4 technical qPCR replicates (from the same cDNA) and took the median value for further analysis.

### Inhibitor treatment and infection assay

We seeded 10^5^ HuH-7 cells per well of a 24-well plate. The next day, growth medium was replaced by medium containing Cyclosporin A (Sigma-Aldrich, SML1018) or BX-795 hydrochloride (Sigma-Aldrich, SML0694) at indicated concentrations 2h prior to infection. Cells were infected with SARS-CoV-2 at MOI 0.5 PFU/cell by adding virus on top of the inhibitor-containing media.

### Cell viability assay

We seeded 2*10^4^ HuH-7 cells per well of a 96-well plate. The next day, the growth medium was replaced by medium containing Cyclosporin A (Sigma-Aldrich, SML1018) or BX-795 hydrochloride (Sigma-Aldrich, SML0694) at indicated concentrations. Cells were incubated for 72 h, and cell viability was assessed using the Cell-Titer-Glo reagent (Promega) according to the manufacturer’s instructions.

### Quantification of ribosomal frameshifting

HEK293 cells were transiently transfected with either the control construct or the frameshifting construct of our dual-color EGFP-mCherry translation reporter outlined in Supplementary Figure 3a. RYDEN was expressed as fusion proteins with ECFP. In control experiments a plasmid only carrying ECFP was used. Using flow cytometry (Novocyte Quanteon), ECFP-positive cells were analyzed for the ratio between mCherry and EGFP, providing a direct readout of ribosomal frameshifting efficiency. Accordingly, frameshifting efficiency was calculated using the ratio of mCherry to EGFP observed with the frameshifting reporter construct (Supplementary Figure 3a), relative to the mCherry/EGFP ratio observed with the control construct (Supplementary Figure 3a).

### Computational analyses

#### Protein-protein interaction network

To establish protein-protein interactions for the proteins identified from the MS experiments, we utilized STRING v11^93^. Specifically, for the RAP-MS network (Figure 2), we seeded all proteins detected with an adj. *P* < 0.2 and positive logFC from the moderated *t*-test between SARS-CoV2 purifications and RMRP purifications. Edges between interacting proteins were included for those above a combined interaction score of 550. To generate the combined RAP-MS and proteome MS network, we seeded nodes where the adj. *P* < 0.05 for either of the assays. Edges between RAP-MS nodes and proteome MS nodes were included for combined interaction scores exceeding 700.

### Gene set and pathway enrichment analysis

We performed a hypergeometric Gene Ontology (GO) enrichment analysis for the expanded SARS-CoV-2 interactome proteins using the Database for Annotation, Visualization and Integrated Discovery (DAVID) tool (https://david.ncifcrf.gov/tools.jsp) and applying default settings.

We performed Gene Set Enrichment Analysis (GSEA) for proteome experiments with the clusterProfilter R package^94^ utilizing the Hallmark and C5-Biological Processes gene sets available through MSigDB^95^. Genes were ranked based on the product of the log2 fold change and the log10 moderated *t*-test *P* value between the SARS-CoV2 treatment and mock treatments.

### eCLIP and RNA sequencing analysis

Paired-end sequencing reads from (i) eCLIP experiments, or (ii) sequencing of crosslinked RNA fragments following RAP-MS, were trimmed using a custom python script that simultaneously identified the umi-molecular identifier (UMI) associated with each read. These trimmed reads were then aligned to the SARS-Cov2 reference genome (NC_045512.2 contig) using bwa^96^. Next, we removed PCR duplicates using the UMI-aware deduplication functionality in Picard’s MarkDuplicates. Finally, enriched regions of binding were identified using the MACS2^97^ to model the fold change between per-million fragment normalized counts (--SPMR) of the treated and control. Visualizations of the region were rendered from the PCR-deduplicated .bam files using the Integrative Genome Visualization (IGV) Browser.

### Code and data availability

Custom computer code and an interactive rendering of our protein-protein association network of SARS-CoV-2 RNA interacting proteins is publicly accessibly at: https://munschauerlab.github.io/SCoV2-proteome-atlas/.

## ACKNOWLEDGMENTS

We thank D.R. Mani for help with statistical analysis; C. Vockley, R. Smyth, A.-E. Saliba, L. Barquist, V. Subramanian, S. Myers, and the Munschauer group for helpful discussions and critical comments on the manuscript; K. Clauser for help with the mass spectrometry database and data searches; L. Gaffney for help with artwork. Work in the Munschauer Laboratory is supported by the Helmholtz Association.

## COMPETING INTERESTS

The authors declare no competing interests.

## Notes

https://munschauerlab.github.io/SCoV2-proteome-atlas/

### Competing Interest Statement

The authors have declared no competing interest.

