## Supplementary Figures for "A direct RNA-protein interaction atlas of the SARS-CoV-2 RNA in infected human cells"

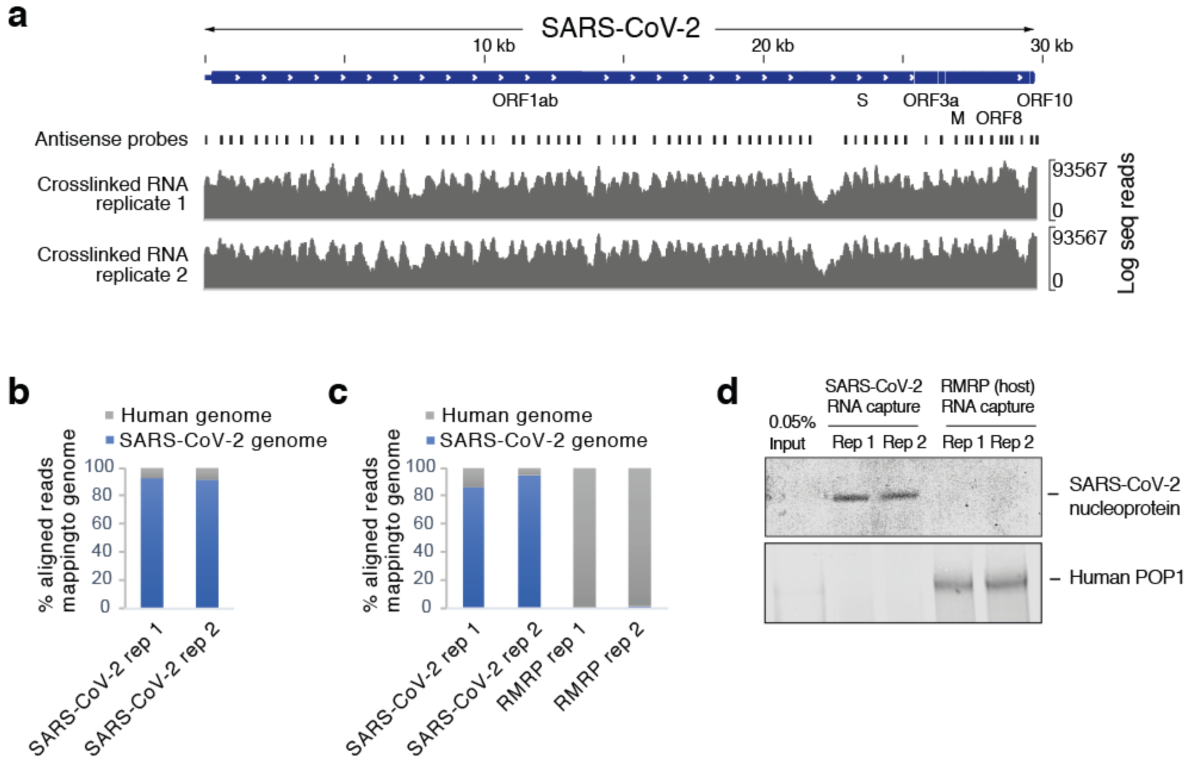


**Supplementary Figure 1 | Capturing the SARS-CoV-2 RNA genome and its bound proteins with RAP-MS. a,** Alignment of protein-crosslinked RNA fragments to the SARS-CoV-2 genome following RNA antisense purification of SARS-CoV-2. Two replicate experiments are shown. **b,** Fraction of all aligned sequence reads mapping to the human or SARS-CoV-2 genomes in pilot RAP-MS experiment. **c**, As in **b**, but for full scale SARS-CoV-2 RAP-MS and RMRP RAP-MS experiments. **d**, Western blot of SARS-CoV-2 and RMRP RAP-MS experiments. Antibodies for SARS-CoV-2 nucleoprotein (Abcam, ab272852) and human POP1 (Proteintech, 12029-1-AP) were used.

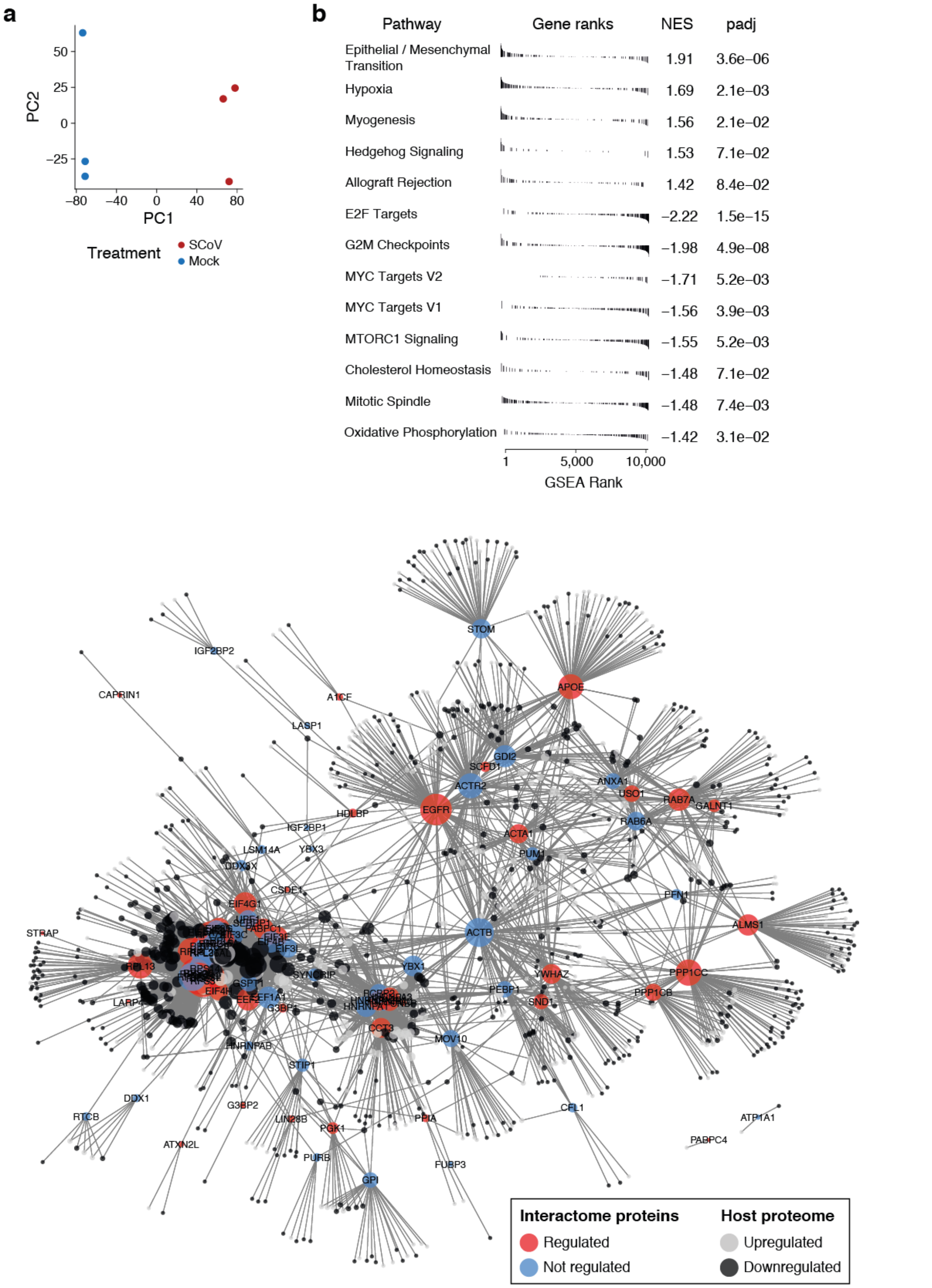


**Supplementary Figure 2 | Connecting the SARS-CoV-2 RNA interactome to proteome dynamics in infected cells. a,** Principle component analysis for proteome measurements of SARS-CoV-2 infected or Mock infected HuH-7 cells. **b,** Gene set enrichment analysis for proteins significantly regulated in global proteome measurements. Gene sets enriched in addition to those shown in Figure 3b are presented. **c,** Protein-protein association network of expanded SARS-CoV-2 interactome proteins (blue: interactome protein, not regulated; red: interactome protein, regulated) and their connections to differentially regulated proteins upon SARS-CoV-2 infection. Upregulated proteins are shown in light grey; downregulated proteins are shown in dark grey. Circle sizes scale to the number of connections of each interactome protein.

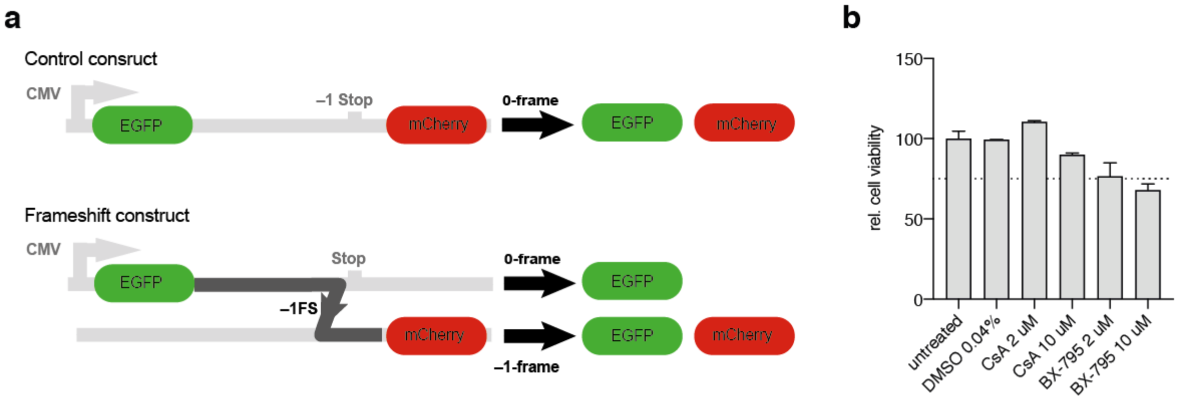


**Supplementary Figure 3 | Functional validation of SARS-CoV-2-binding proteins. a,** Schematic of dual-fluorescence translation reporter to quantify ribosomal frameshifting efficiency. The depicted control construct contains EGFP and mCherry in an in-frame orientation, leading to the production of both fluorescence proteins separated by a self-cleaving 2A peptide when the 0 reading frame is translated. In the frameshift construct depicted below, EGFP and mCherry are separated by an in-frame stop codon, preventing the production of mCherry when the 0 reading frame is translated. −1FS leads to the production of EGFP and mCherry and the ratio between both fluorescence proteins is a direct measure of frameshifting efficacy. –1FS: –1 ribosomal frameshifting. **b**, Cell viability assay in inhibitor-treated and untreated HuH-7 cells (*n*=2).

### SUPPLEMENTARY TABLES

**Supplementary Table 1 |** Proteins detected by quantitative mass spectrometry in SARS-CoV-2 and RMRP RNA antisense purifications in infected human cells.

**Supplementary Table 2 |** Intersection of the expanded SARS-CoV-2 RNA interactome with published work.

**Supplementary Table 3 |** Gene Ontology enrichment analysis for the expanded SARS-CoV-2 RNA interactome.

**Supplementary Table 4 |** Protein-protein association network based on STRING v11 interactions between human proteins in the expanded SARS-CoV-2 RNA interactome.

**Supplementary Table 5 |** Proteome abundance measurements in SARS-CoV-2 infected and uninfected HuH-7 cells.

**Supplementary Table 6 |** Gene set enrichment analysis for proteome abundance measurements in SARS-CoV-2 infected cells HuH-7 cells.

**Supplementary Table 7 |** Protein-protein association network based on STRING v11 interactions between human proteins in the core SARS-CoV-2 RNA interactome and proteins regulated upon SARS-CoV-2 infection in HuH-7 cells.
